# A systematic evaluation of IgM and IgG antibody assay accuracy in diagnosing acute Zika Virus infection in Brazil; lessons relevant to emerging infections

**DOI:** 10.1101/2020.11.25.399386

**Authors:** Raquel Medialdea-Carrera, Flavia Levy, Priscila Castanha, Patricia Carvalho de Sequeira, Patricia Brasil, Lia L Lewis-Ximenez, Lance Turtle, Tom Solomon, Ana Maria Bispo de Filippis, David W. Brown, Michael J. Griffiths

**Author notes:** These authors contributed equally to the work (David W. Brown, Michael J. Griffiths). Address for correspondence: Department of Clinical Infection, Microbiology and Immunology, Ronald Ross Building, 8 West Derby Street, Institute of Infection and Global Health, University of Liverpool, L69 7BE, UK.

## Abstract

Accurate diagnostics underpin effective public health responses to emerging viruses. For viruses, such as Zika virus (ZIKV), where the viremia clears quickly, antibody-based (IgM or IgG) diagnostics are recommended for patients who present seven days after symptom onset. However, cross-reactive antibody responses can complicate test interpretation among populations where closely related viruses circulate.

We examined the accuracy (proportion of samples correctly categorized as Zika-positive or negative) for antibody-based diagnostics among Brazilian residents (Rio de Janeiro) during the ZIKV outbreak. Four ZIKV ELISAs (IgM and IgG Euroimmun, IgM Novagnost and CDC MAC), two dengue ELISAs (IgM and IgG Panbio), and the ZIKV plaque reduction neutralization test (PRNT) were evaluated. Positive samples were ZIKV PCR confirmed clinical cases collected in 2015-2016 (n=169); Negative samples (n=236) were collected before ZIKV was present in Brazil (≤2013).

Among serum samples collected ≥7 days from symptom onset, PRNT exhibited the highest accuracy (93.7%), followed by the Euroimmun IgG ELISA (77.9%). All IgM assays exhibited lower accuracy (<74%). IgG was detected more consistently than IgM among ZIKV cases using Euroimmun ELISAs (68% versus 22%). Anti-DENV IgM ELISA was positive in 41.1% of confirmed ZIKV samples tested.

The Euroimmun IgG assay, although misdiagnosing 22% of samples, provided the most accurate ELISA. Anti-ZIKV IgG was detected more reliably than IgM among ZIKV patients, suggesting a secondary antibody response to assay antigens following ZIKV infection. Antibody ELISAs need careful evaluation in their target population to optimise use and minimise misdiagnosis, prior to widespread deployment, particularly where related viruses co-circulate.

## Introduction

Zika virus (ZIKV) is an arthropod-borne flavivirus. A public health emergency of international concern (PHEIC) was declared by the World Health Organization (WHO), in response to the large Zika epidemics in South and Central America in 2015-2016 (1). Although transmission has declined, over 85 countries across South and Central America, Asia, West Africa, Caribbean and the Pacific Islands have current or previous transmission of ZIKV and another 61 have stablished mosquito vectors and remain at risk for Zika infection (2–4). Moreover, autochthonous transmission of Zika Virus was demonstrated in Europe with, at least, 3 cases of vector-borne transmission in Southern France in 2019 (5). Accurate diagnostics are essential to guide appropriate clinical management of suspected patients. Both false negative and false positive diagnosis may trigger catastrophic consequences, especially among pregnant women (6).

The first-line diagnostic test for ZIKV detection is direct molecular detection using PCR. However, the time-frame for accurate virus detection following exposure using this method is short. Consequently, the WHO issued laboratory diagnostic algorithms recommending anti-ZIKV antibody-based testing in patients presenting seven or more days after symptom onset (7, 8). Nevertheless, accurate detection of ZIKV antibodies is challenging because antibody-based assays are susceptible to cross-reactivity from related viruses. This is a particular issue in regions such as Latin America where extensive circulation of multiple flaviviruses has occurred in the population over the last 30 years. In Brazil, all four serotypes of dengue virus (DENV) circulate and yellow fever virus (YFV) vaccination is widespread in many regions (9). Studies have shown that over 53% of pregnant women are reported as anti-DENV immunoglobulin G (IgG) positive in Brazil and in certain regions, dengue seroprevalence is estimated to be over 75% (10, 11). Similarly, yellow fever vaccination has been reported to have reasonable coverage in most regions of Brazil and coverage is likely to have increased following the vaccination campaigns mounted in response to recent yellow fever outbreaks (12, 13).

The WHO highlighted the important need for field validation of available Zika serological assays in flavivirus exposed populations (14). A range of different antibody-based assays have been developed (15, 16), but evaluation of the assay’s performance in local populations lagged behind their use.

To date, there are very few published studies on the performance of commercial ZIKV antibody-based assays using well-characterized samples from South American populations and no systematic evaluation from Brazil. Evaluations based on sera from European travelers who had visited countries with ZIKV circulation found a high sensitivity and specificity for the commercial immunoglobulin M (IgM) and IgG enzyme-linked immunosorbent assays (ELISAs) that use a recombinant ZIKV NS1 antigen (17–20). Other commercial assays, such as the IgM μ-capture ELISA, have also been approved for use in the Americas, despite initial reports of low test specificity among travelers (21). The Center for Disease Control and Prevention (CDC) in the USA, recommends the use of an IgM antibody capture enzyme-linked immunosorbent assay (MAC-ELISA) which is licensed under the CDC FDA-emergency-use-authorization protocol. However, reports from Nicaragua and the USA have shown this assay to have relatively low specificity and it is no longer recommended for screening. Further local population evaluations have been recommended (22, 23).

We conducted a systematic evaluation of four antibody-based methods for ZIKV diagnosis: the IgM and IgG NS1 Anti-ZIKV ELISAs (Euroimmun), the IgM μ-capture ZIKV ELISA (Novagnost) and the CDC MAC-ELISA. We also re-evaluated two DENV antibody assays, the IgM and IgG ELISA assays (Panbio) in this new context. We compared these ELISAs against the ZIKV plaque reduction neutralization test (PRNT), which is currently considered the “gold standard” by the WHO for the confirmatory diagnosis of Flavivirus infections. The assays were evaluated using well-characterized sera from residents of Rio de Janeiro, Brazil, classified by clinical and laboratory testing, as confirmed ZIKV (cases with both clinical evidence of ZIKV infection and positive detection of ZIKV RNA by RT-PCR in at least one specimen) or non-ZIKV cases (sera collected in 2013 or before, prior to the arrival of ZIKV in Rio de Janeiro). Our aim was to examine assay accuracy among the Brazilian population, to investigate the time window of detection of anti-ZIKV antibodies and the biological variability of antibody responses.

## Materials and Methods

### Ethics Statement

The sera and patient data were used in accordance with the ethical standards of the Instituto Nacional de Infectologia Evandro Chagas and the Instituto Oswaldo Cruz. The study protocol was approved by its Research Ethics Committee (reference CAAE 0026.0.009.000–07 and CAAE 71405717.8.0000.5248). Specimens were given laboratory numbers and so anonymized to testers to ensure patient confidentiality.

### Study population and sample selection

ELISA evaluations were performed at the Flavivirus Reference Laboratory, Institute Oswaldo Cruz in Rio de Janeiro (Brazil). Plaque reduction neutralization tests (PRNT) were performed at the Aggeu Magalhães Institute in Recife (Brazil).

Four-hundred and five serum samples in total were tested (Table 1). All sera were collected from residents (n=307) of the State of Rio de Janeiro. Only samples stored at −20°C or below, and with no history of repeated freeze-thaw were used in the study. Due to limited availability of reagents and serum volumes, not all samples were tested on all assays.

**Table 1.** Characteristics of the study population

|  |  | No. of patients | No. of samples | % of patients | Year. |
| --- | --- | --- | --- | --- | --- |
| **Total** |  | 307 | 405 | - | 2002-2016 |
| **Sex** | Female | 164 (296)* | - | 55.4% | - |
| **ZIKV Positive samples (Set1)** |  |  |  |  |  |
|  | **Total** | **71** | **169** | **23.1 (71/307)<sup>a</sup>** | **2015-2016** |
|  | 1 Serum | 5 | 5 | 7.0 (5/71)<sup>b</sup> | 2015-2016 |
|  | 2 Serum | 55 | 110 | 77.5(55/71)<sup>b</sup> | 2015-2016 |
|  | 3 or more samples | 11 | 54 | 15.5(11/71)<sup>b</sup> | 2015-2016 |
| **Controls – ZIKV Negative (Set2)** |  |  |  |  |  |
|  | **Total** | **184** | **184** | **59.9(184/307)<sup>a</sup>** | **2002-2013** |
|  | DENV1-4 (Total) | 90 | 90 | 82.6 (90/109)<sup>d</sup> | 2002-2013 |
|  | DENV 1 | 21 | 21 | 23.3(21/90)<sup>c</sup> | 2010-2011 |
| **DENV** | DENV 2 | 17 | 17 | 18.9(17/90)<sup>c</sup> | 2008,2010,2011 |
|  | DENV 3 | 21 | 21 | 23.3(21/90)<sup>c</sup> | 2002,2007,2008 |
|  | DENV 4 | 31 | 31 | 34.4(31/90)<sup>c</sup> | 2012,2013 |
| **Yellow fever** |  | 19 | 19 | 17.4(19/109)<sup>d</sup> | 2003-2007 |
| **Measles or Rubella** |  | 40 | 40 | 21.7(40/184)<sup>e</sup> | 2011-2012 |
| **Other non-flavivirus infections** |  | 35 | 35 | 19.0(35/184)<sup>e</sup> | 2011-2012 |
| **General population (Set3)** | Total Set3 | 52 | 52 | 16.9(52/307) a | 2013 |

Abbreviations: No., number, Year, Year of sample collection, \*Sex was not documented for 18 patients.
The population data for the subgroups denoted by roman superscript letters a through e add up to 100%:
<sup>a</sup>. represents patients in Set1, 2 and 3. <sup>b</sup>. Patients in Set1. <sup>c</sup>. Dengue Positive patients. <sup>d</sup>. Flavivirus positive subjects. <sup>e</sup>. Non-flavivirus infections subjects.

Panels of sera were assembled for the evaluation: 1) to assess assay sensitivity, samples from ZIKV PCR confirmed cases were used (Set1: n=169) from subjects (n=71) with rash-fever symptom and positive detection of ZIKV RNA by RT-PCR. Samples were collected in 2015 or 2016, coinciding with the peak of the ZIKV epidemic in Rio de Janeiro (1). Most subjects (66 out of 71) had two or more sequential samples collected (see Table S1 for further details on serum collections). 2) To assess assay specificity, non-ZIKV samples, collected from individuals during, or before, 2013 were used (Set2 and Set3; n=236). These were collected 2 years before the ZIKV outbreak in Rio de Janeiro began in 2015 (24). Set2 (n=184) included samples from subjects with confirmed dengue (clinical presentation and PCR or IgM positive; n=90), measles or rubella infection (n=40), who had received yellow fever vaccination (n=19), or other population samples (n=35). Set3 samples were collected from patients attending a hepatitis clinic at Fiocruz, Rio de Janeiro (n=52).

### Diagnostic Assays

The IgM and IgG ZIKV NS1 ELISA assays (Euroimmun, Lubeck, Germany) use recombinant Zika non-structural protein 1(NS1) as the ZIKV antigen. Assays were performed following the manufacturer’s instructions. IgM and IgG results were determined based on optical density (OD) ratio of human sample/calibrator sample. A result was classified as: positive if ratio ≥1.1; indeterminate between ≥0.8 and 1.1; negative ≤0.8.

The Novagnost Zika Virus IgM μ-capture ELISA (NovaTec Immunodiagnostica GmbH, Germany) uses ZIKV NS1 antigen. Assays were conducted according to the manufacturer’s instructions. Results were classified based on the OD ratio of human/calibrator sample as per manufacturer’s guidelines.

The CDC ZIKV MAC-ELISA (FDA CDC-designed IgM antibody capture ELISA) utilizes inactivated whole virus antigen. The assays were performed as previously described using the US Centers for Disease Control and Prevention (CDC) emergency use authorization protocol (25). Results were based on the Positive to Negative ratio (P/N). P is obtained as the mean OD of the test serum, which is compared with N, the mean OD of the normal human serum/OD negative-control serum. Results were reported as recommended: positive if P/N≥3; indeterminate if P/N≥2 but 3 and negative if P/N2.

For the detection of DENV IgM and IgG antibodies, the commercial dengue IgG Indirect and IgM Capture ELISAs (Panbio, Alere, United Kingdom) were used. Samples were considered positive for previous or recent dengue infection according to the standard protocols of the manufacturer.

All the ELISA assays used the same volume of patient serum. All assays were stored at 2-8°C prior to use.

PRNTs were performed following a published method (26), it is an adaptation of an established PRNT protocol (27, 28). PRNT was performed using a Zika virus strain isolated in Northeast Brazil (ZIKV, BR-PE243/2015). The cut-off for PRNT positivity was defined based on a 50% reduction in plaque counts (PRNT50). ZIKV neutralizing antibody titers were estimated using a four-parameter non-linear regression. Serum samples were considered positive when antibody titers were >1:100 (log2).

### RT-PCR

Individuals with clinical presentation consistent with acute ZIKV infection (during 2015 and 2016) were tested for detection of ZIKV nucleic acid material. RNA was extracted using the Qiamp Mini Elute (Qiagen, Brazil). Reference ZIKV and DENV RT-PCR (for DENV suspected samples) were performed as previously described (29, 30).

### Statistical analyses and calculation of ROC Curves, sensitivity and specificity

Diagnostic performance of the ELISAs was assessed by calculating accuracy (classified as true positive and true negative / all cases tested); sensitivity (classified as positive / all true positives tested) and specificity (classified as negative / all true negatives tested), using MedCalc Statistical Software, version 16.2.0 (MedCalc Software, Ostend, Belgium). Figures were generated using STATA v16 and PRISM GraphPad v7 (GraphPad Software, La Jolla, CA). Samples with indeterminate or borderline results were re-tested (if enough specimen was available). Samples with indeterminate results a second time were considered negative.

This prospective cross-sectional diagnostic accuracy study is reported according to the Standards for Reporting of Diagnostic Accuracy Study (STARD) (31) (Checklist S1, Flow chart S1, Flow chart S2).

## Results

Age distribution was unimodal with a median age of 24.5 years (range 1-80y). 55.4% of confirmed ZIKV and non-ZIKV cases were female. Among the confirmed ZIKV samples, the median time of sample collection after symptom onset was 7 days (range 1 - 276). Full details are given in Table 1.

### Diagnostic assays performance

A detailed breakdown of accuracy, sensitivity and specificity for all assays tested are presented in Table 2 and Table 3.

**Table 2.**
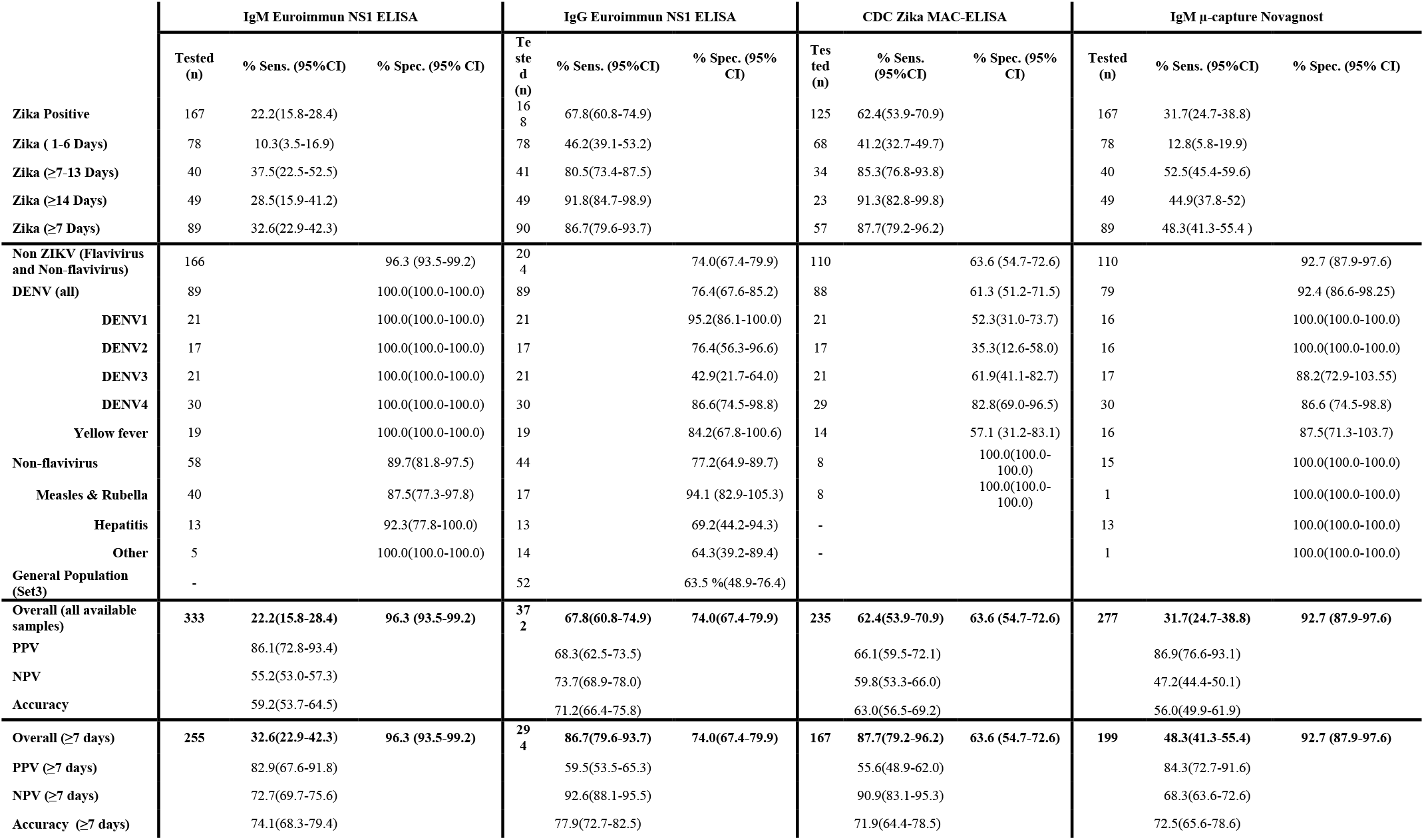
Sensitivity and specificity for the four anti-ZIKV antibody ELISAs. Sensitivity and specificity of the IgM and IgG Euroimmun NS1 commercial ELISA, the MAC-ELISA and IgM μ-capture Novagnost assays with sera from confirmed ZIKV (Set 1) and the Control non-ZIKV group (Set 1 and Set 2). Specificity values were calculated for each assay based on Set 2 and 3. Data from ZIKV-positive cases served only for determining the sensitivity and was not used for the specificity calculation. Similarly, data from ZIKV-negative cases serve only for determining the specificity and were not used for calculating the sensitivity. Overall sensitivity, PPV, NPV and accuracy were calculated with both a) all the ZIKV Positive samples and b) only the ZIKV samples collected ≥7 days post symptom onset. Note: Sens., Sensitivity; Spec., Specificity; CI, Confidence Interval; PPV, Positive Predictive Value; NPV, Negative Predictive Value; ZIKV, Zika virus; DENV, Dengue virus; Days, number of days the sample was collected after symptom onset. Indeterminate results were considered negative for the calculation. Non-flavivirus includes measles & rubella, hepatitis and general population samples. DENV (all) includes all DENV samples (DENV 1-4).

| | IgM Euroimmun NS1 ELISA | | | IgG Euroimmun NS1 ELISA | | | CDC Zika MAC-ELISA | | | IgM μ-capture Novagnost | | |
| --- | --- | --- | --- | --- | --- | --- | --- | --- | --- | --- | --- | --- |
|  | Tested (n) | % Sens. (95%CI) | % Spec. (95% CI) | Tested (n) | % Sens. (95%CI) | % Spec. (95% CI) | Tested (n) | % Sens. (95%CI) | % Spec. (95% CI) | Tested (n) | % Sens. (95%CI) | % Spec. (95% CI) |
| **Zika Positive** | 167 | 22.2(15.8-28.4) |  | 168 | 67.8(60.8-74.9) |  | 125 | 62.4(53.9-70.9) |  | 167 | 31.7(24.7-38.8) |  |
| **Zika ( 1-6 Days)** | 78 | 10.3(3.5-16.9) |  | 78 | 46.2(39.1-53.2) |  | 68 | 41.2(32.7-49.7) |  | 78 | 12.8(5.8-19.9) |  |
| **Zika (≥7-13 Days)** | 40 | 37.5(22.5-52.5) |  | 41 | 80.5(73.4-87.5) |  | 34 | 85.3(76.8-93.8) |  | 40 | 52.5(45.4-59.6) |  |
| **Zika (≥14 Days)** | 49 | 28.5(15.9-41.2) |  | 49 | 91.8(84.7-98.9) |  | 23 | 91.3(82.8-99.8) |  | 49 | 44.9(37.8-52) |  |
| **Zika (≥7 Days)** | 89 | 32.6(22.9-42.3) |  | 90 | 86.7(79.6-93.7) |  | 57 | 87.7(79.2-96.2) |  | 89 | 48.3(41.3-55.4 ) |  |
| **Non ZIKV (Flavivirus and Non-flavivirus)** | 166 |  | 96.3 (93.5-99.2) | 204 |  | 74.0(67.4-79.9) | 110 |  | 63.6 (54.7-72.6) | 110 |  | 92.7 (87.9-97.6) |
| **DENV (all)** | 89 |  | 100.0(100.0-100.0) | 89 |  | 76.4(67.6-85.2) | 88 |  | 61.3 (51.2-71.5) | 79 |  | 92.4 (86.6-98.25) |
| **DENV1** | 21 |  | 100.0(100.0-100.0) | 21 |  | 95.2(86.1-100.0) | 21 |  | 52.3(31.0-73.7) | 16 |  | 100.0(100.0-100.0) |
| **DENV2** | 17 |  | 100.0(100.0-100.0) | 17 |  | 76.4(56.3-96.6) | 17 |  | 35.3(12.6-58.0) | 16 |  | 100.0(100.0-100.0) |
| **DENV3** | 21 |  | 100.0(100.0-100.0) | 21 |  | 42.9(21.7-64.0) | 21 |  | 61.9(41.1-82.7) | 17 |  | 88.2(72.9-103.55) |
| **DENV4** | 30 |  | 100.0(100.0-100.0) | 30 |  | 86.6(74.5-98.8) | 29 |  | 82.8(69.0-96.5) | 30 |  | 86.6 (74.5-98.8) |
| **Yellow fever** | 19 |  | 100.0(100.0-100.0) | 19 |  | 84.2(67.8-100.6) | 14 |  | 57.1 (31.2-83.1) | 16 |  | 87.5(71.3-103.7) |
| **Non-flavivirus** | 58 |  | 89.7(81.8-97.5) | 44 |  | 77.2(64.9-89.7) | 8 |  | 100.0(100.0-100.0) | 15 |  | 100.0(100.0-100.0) |
| **Measles & Rubella** | 40 |  | 87.5(77.3-97.8) | 17 |  | 94.1 (82.9-105.3) | 8 |  | 100.0(100.0-100.0) | 1 |  | 100.0(100.0-100.0) |
| **Hepatitis** | 13 |  | 92.3(77.8-100.0) | 13 |  | 69.2(44.2-94.3) | - |  |  | 13 |  | 100.0(100.0-100.0) |
| **Other** | 5 |  | 100.0(100.0-100.0) | 14 |  | 64.3(39.2-89.4) | - |  |  | 1 |  | 100.0(100.0-100.0) |
| **General Population (Set3)** | - |  |  | 52 |  | 63.5 %(48.9-76.4) |  |  |  |  |  |  |
| **Overall (all available samples)** | 333 | 22.2(15.8-28.4) | 96.3 (93.5-99.2) | 372 | 67.8(60.8-74.9) | 74.0(67.4-79.9) | 235 | 62.4(53.9-70.9) | 63.6 (54.7-72.6) | 277 | 31.7(24.7-38.8) | 92.7 (87.9-97.6) |
| **PPV** |  | 86.1(72.8-93.4) |  |  | 68.3(62.5-73.5) |  |  | 66.1(59.5-72.1) |  |  | 86.9(76.6-93.1) |  |
| **NPV** |  | 55.2(53.0-57.3) |  |  | 73.7(68.9-78.0) |  |  | 59.8(53.3-66.0) |  |  | 47.2(44.4-50.1) |  |
| **Accuracy** |  | 59.2(53.7-64.5) |  |  | 71.2(66.4-75.8) |  |  | 63.0(56.5-69.2) |  |  | 56.0(49.9-61.9) |  |
| **Overall (≥7 days)** | 255 | 32.6(22.9-42.3) | 96.3 (93.5-99.2) | 294 | 86.7(79.6-93.7) | 74.0(67.4-79.9) | 167 | 87.7(79.2-96.2) | 63.6 (54.7-72.6) | 199 | 48.3(41.3-55.4) | 92.7 (87.9-97.6) |
| **PPV (≥7 days)** |  | 82.9(67.6-91.8) |  |  | 59.5(53.5-65.3) |  |  | 55.6(48.9-62.0) |  |  | 84.3(72.7-91.6) |  |
| **NPV (≥7 days)** |  | 72.7(69.7-75.6) |  |  | 92.6(88.1-95.5) |  |  | 90.9(83.1-95.3) |  |  | 68.3(63.6-72.6) |  |
| **Accuracy (≥7 days)** |  | 74.1(68.3-79.4) |  |  | 77.9(72.7-82.5) |  |  | 71.9(64.4-78.5) |  |  | 72.5(65.6-78.6) |  |

**Table 3.**
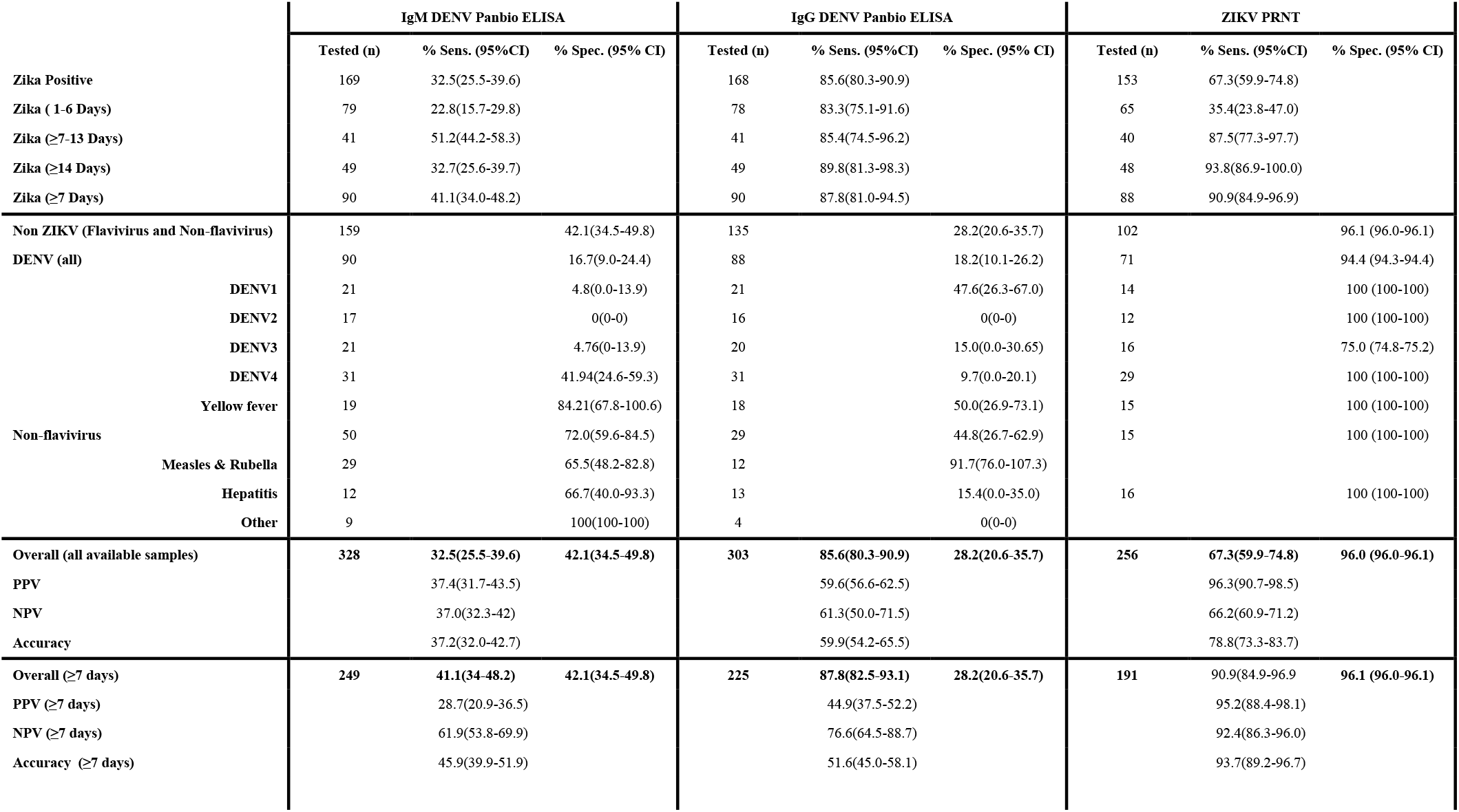
Sensitivity and specificity for the two anti-DENV antibody ELISAs and the ZIKV PRNT. The sensitivity and specificity of the IgM and IgG Panbio DENV commercial ELISA and ZIKV PRNT with the ZIKV panel (Set 1) and the Control non-ZIKV group (Set 2). Specificity values were calculated for each assay based on Set 2. Data from ZIKV-positive cases served only for determining the sensitivity and was not used for the specificity calculation. Similarly, data from ZIKV-negative cases serve only for determining the specificity and were not used for calculating the sensitivity. Overall sensitivity, PPV, NPV and accuracy were calculated with ZIKV Positive samples collected ≥7 days post symptom onset. Note: Sens., Sensitivity; Spec., Specificity; CI, Coefficient Interval; PPV, Positive Predictive Value; NPV, Negative Predictive Value; ZIKV, Zika virus; DENV, Dengue virus; Days, number of days the sample was collected after symptom onset. Indeterminate results were considered negative for the calculation. Non-flavivirus includes measles & rubella, hepatitis and general population samples. DENV (all) includes all DENV samples (DENV 1-4).

### Summary of overall accuracy

Initially, we examined test accuracy among all samples, including samples collected within 7 days of ZIKV symptom onset. The highest accuracy was exhibited by PRNT, followed by ZIKV IgG NS1 Euroimmun, MAC-ELISA, IgM NS1 Euroimmun, and IgM μ-capture Novagnost ELISAs (78.8, 71.2, 63.6, 59.2 and 56.0% respectively).

We also assessed the performance of the dengue assays (IgG and IgM) for detecting ZIKV antibodies. The DENV IgG ELISA exhibited similar results to the ZIKV IgG assays (59.7% [181/303]) of samples correctly classified). The DENV IgM ELISA demonstrated 37.2% (122/328) samples correctly classified for ZIKV.

The WHO recommends that samples collected up to 7 days from symptom onset are tested using a ZIKV specific RT-PCR (32). Analysis was repeated excluding ZIKV samples collected less than 7 days from symptom onset. The accuracy of all ZIKV assays improved. Again, the highest accuracy was exhibited by PRNT followed by the IgG NS1 Euroimmun ELISA (93.7 and 77.9 respectively). The ZIKV IgM (NS1 Euroimmun, μ-capture Novagnost and CDC MAC) ELISAs continued to exhibit lower accuracy (73.6, 72.5 and 71.9% respectively).

PRNT exhibited both good sensitivity (89.8% [79/88]) and specificity (96.1% [98/102]). The major weakness of the Euroimmun IgM and Novagnost IgM ELISAs was poor sensitivity (32.6 [29/89] and 48.3% [43/89] respectively). The CDC MAC-ELISA exhibited reasonable sensitivity but poor specificity (87.7% [50/57] and 63.6% [70/110] respectively; Tables 2 and 3). Assay sensitivity improved in all the assays among samples collected ≥7 to 13 days post symptom onset. Assay sensitivity decreased for specimens collected ≥14 days in the IgM and IgG ZIKV Euroimmun, IgM Novagnost and IgM DENV assay.

### Comparison of anti-ZIKV IgG and IgM antibody responses

We compared IgG and IgM anti-ZIKV antibody titers over time from symptom onset (measured using the ZIKV NS1 Euroimmun ELISAs). Based on measuring a single sample, anti-ZIKV IgG was more consistently detected than anti-ZIKV IgM among sera from confirmed ZIKV positive subjects (68% [114/168] vs. 22% [37/167] respectively). Among paired samples, anti-ZIKV IgG more frequently exhibited a rise in antibody titers than IgM (IgG; rise 95% [56/59]; ≥2 fold 75% [44/59]; IgM; rise 81% [46/57]; ≥2 fold 61% [35/57]; Figure 1). Anti-ZIKV IgG also exhibited a higher median fold increase compared to IgM among paired sera (3.5 versus 2.6-fold increase [p=0.001]; taken 7 days apart [median]; Figure 2). Overall, there was a more sustained rise in anti-ZIKV IgG compared to IgM over time from symptom onset (Figure 3).

**Figure 1.**
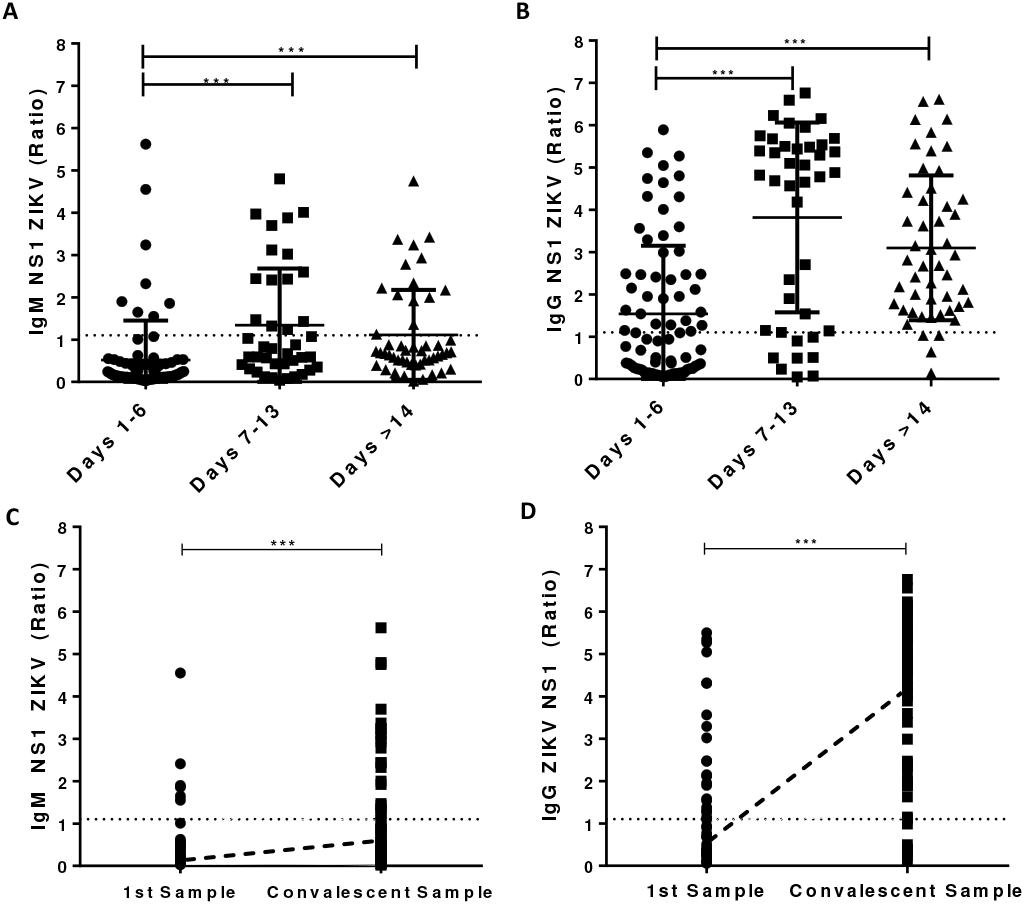
Zika antibody detected in serum samples collected during the acute (1-6 days after onset), early-convalescent phase (7-13d) and late convalescent-phase (≥14d) from PCR positive Zika cases measured by IgM (A) or IgG (B) NS1 anti-ZIKV ELISAs (Euroimmun). C) IgM NS1 anti-ZIKV ELISA measurements for acute (1-6 days after onset) and convalescent (≥7 days) samples from PCR positive ZIKV cases. D) IgG NS1 anti-ZIKV antibody levels in paired serum samples from PCR positive ZIKV positive cases. Dotted horizontal lines represent the cut-off value used in each assay. Data points above the cut-off are considered positive. Trend-line in C) and D) represent the median antibody levels for acute and convalescent samples. Statistically significant differences between two groups were measured by Mann Whitney U test (*** p=0.0001). Figure shows antibody Ratios* calculated as per manufacturers’ instructions (Antibody Ratio = OD Sample/OD Calibrator).

**Figure 2.**
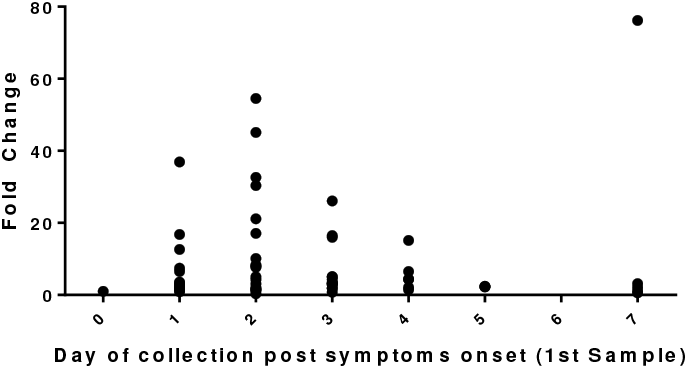
The change in IgG NS1 Euroimmun anti-ZIKV antibody levels between paired serum samples from PCR positive Zika cases by day of collection (days post symptom onset) of the first (acute) sample. Based on the first sample (collected 0 −7 days) and second sample (median interval between samples was 7 days). The highest fold change (change in antibody level between 1^st^ and 2^nd^ samples) was observed among paired samples collected on days 2 and 7 post symptom onset.

**Figure 3.**
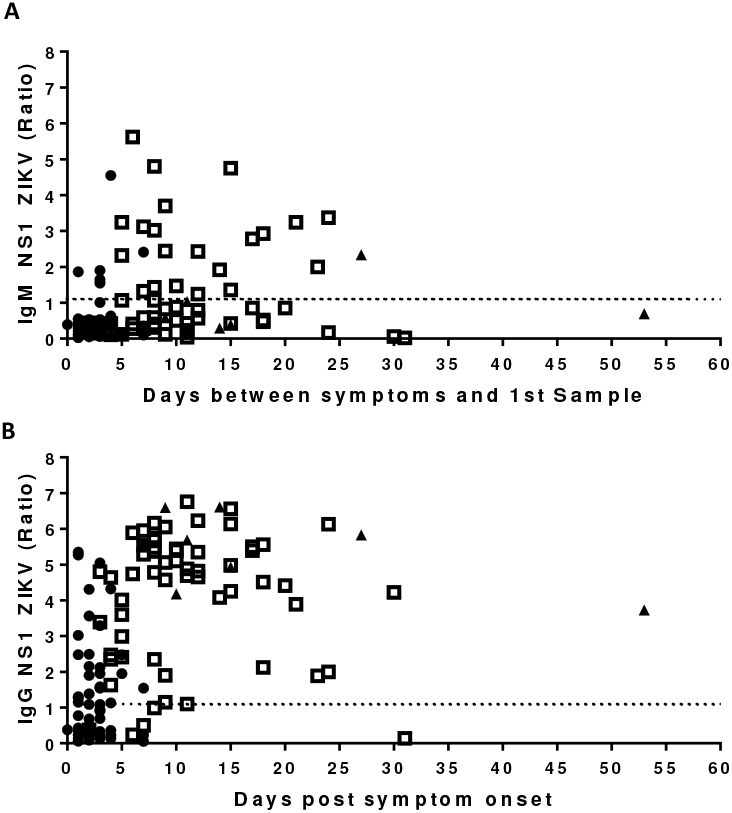
Anti-ZIKV antibody levels in sequential serum samples collected from Zika PCR positive cases on different days post symptom onset (0-54 days) measured in A) IgM Euroimmun NS1 and B) IgG Euroimmun NS1 anti-ZIKV ELISAs. Dotted line shows assay cut-offs. The figure shows more consistent detection of anti-Zika antibodies (level above the cut-off) among convalescent samples when measuring IgG compared to IgM. Ratios calculated as per manufacturers’ instruction; 1st collection (acute sample [closed circles]); 2nd collection (convalescent samples [open squares]); 3rd collection (late convalescent samples [closed triangles])

### Anti-DENV IgM and IgG responses in confirmed ZIKV cases

Both IgM and IgG anti-DENV ELISAs gave positive results among sera collected ≥7 days from PCR confirmed ZIKV cases (41% [37/90] and 88% [79/90] respectively - for dengue IgM these are likely to represent false positive IgM results) (Table 3). In an attempt to distinguish between the dengue IgG response in ZIKV patients who had previously been exposed to DENV and acute ZIKV patients exhibiting a false positive cross-reaction against DENV IgG antibodies, we looked at sera collected < 7 days from symptom onset. A high proportion of confirmed ZIKV cases were DENV IgG positive (83% [65/78]). A significant correlation between anti-ZIKV and anti-DENV antibody titers were observed when the same sera were measured by IgG NS1 Euroimmun ZIKV and IgG Panbio DENV assays (IgG r^2^=0.258; p<0.0001; Figure 4a). A similar, but less significant correlation, was observed between ZIKV Euroimmun IgM NS1 and Panbio DENV IgM assays (IgM r^2^=0.015; p=0.03; Figure 4b).

**Figure 4.**
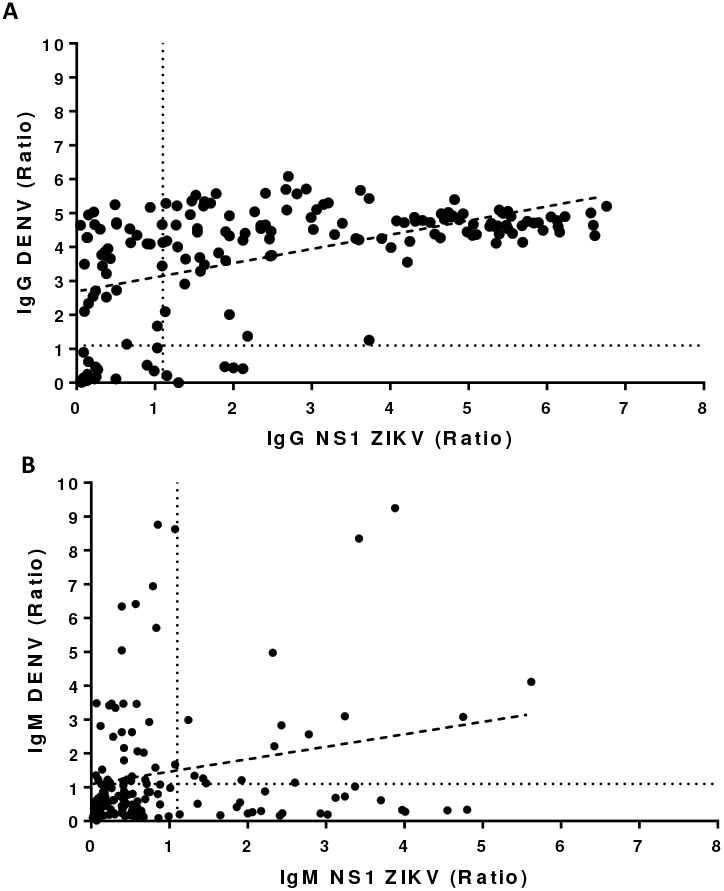
Correlation between anti-ZIKV and anti-DENV antibody levels in individual sera samples. A) IgG anti-DENV ELISA (Panbio) versus IgG anti-ZIKV NS1 (Euroimmun) ELISA. Anti-DENV and anti-ZIKV IgG antibody levels showed a positive correlation. When a patient exhibited a relatively high anti-DENV IgG antibody response they also tended to exhibit a relatively high anti-ZIKV IgG antibody response (p<0.001; r2=0.258; n=168); B) IgM DENV ELISA and IgM ZIKV NS1 ELISA antibody levels. Again the measurements showed a positive correlation (p=0.015; r2=0.015; n=166). Dotted lines show assay cut-offs. Dashed line shows the best-fitting line (Spearman rank correlation [r2]). Correlation was more significant between anti-ZIKV and anti-DENV IgG antibody measurements. The correlation in antibody measurement between ZIKV and DENV ELISAs suggests a degree of overlap in patient responses and/or cross-reaction in antibody detection.

### Diversity of anti-ZIKV IgM antibody responses

To describe the variation in duration and magnitude of anti-ZIKV antibody responses, we compared Zika antibody patterns in four PCR confirmed ZIKV patients (A-D, who had sera collected at ≥5 different time-points after symptom onset (range: 0-276 days). Antibody titers were measured by the IgM NS1 and μ-capture assays (Figure 5). Antibody patterns were diverse in both magnitude and duration of response. Interestingly, in patient A, despite an initial rise in IgM titers, this quickly fell. IgM was below the Euroimmun NS1 ZIKV assay’s positive threshold by day 22. We then looked at all samples from confirmed ZIKV subjects. At 14, 27 and 90 days post symptom onset only 28.6% (14/49), 8% (4/27), and 0% (0/14) samples had detectible IgM responses when measured in the ZIKV Euroimmun NS1 IgM assay. Similarly, only 44.9% (22/49), 25.9% (7/27) and 7.1% (1/14) samples were positive, at 14, 27 and 90 days, when the same samples were measured using the IgM μ-capture Novagnost assay.

**Figure 5.**
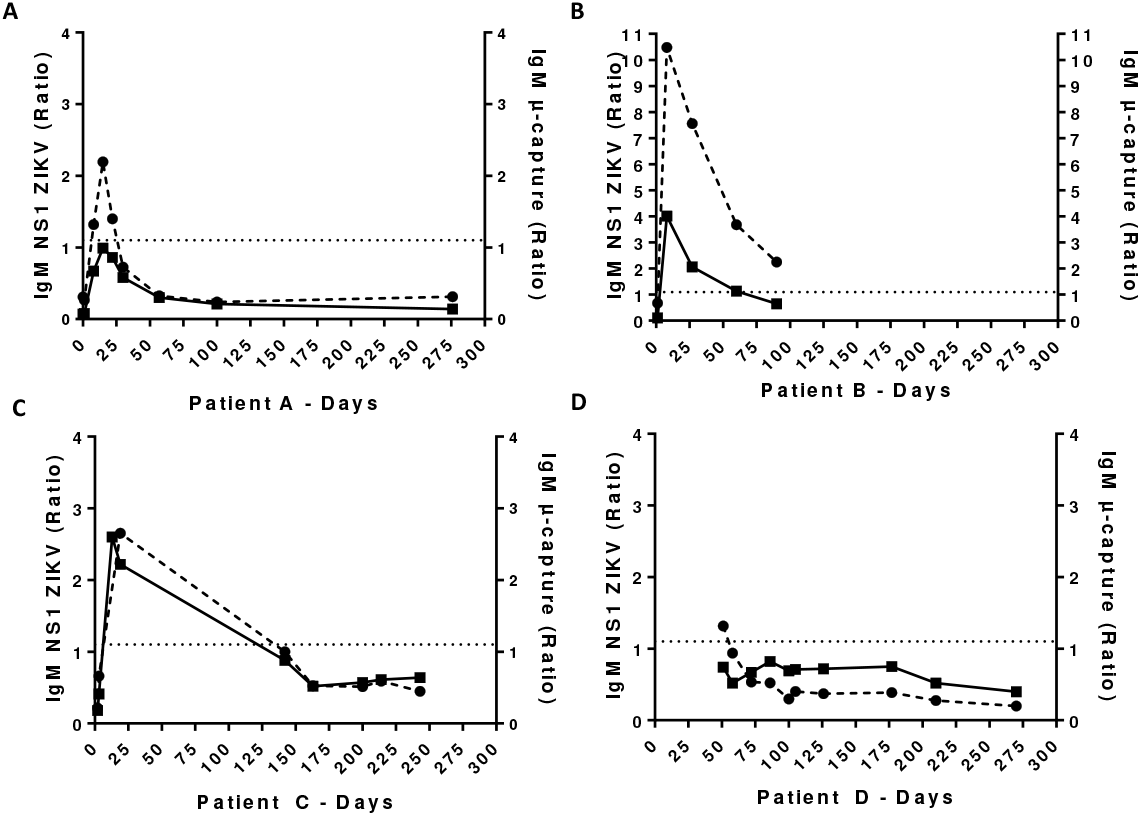
The change in anti-ZIKV IgM antibody levels by day post symptom onset in sequential sera from ZIKV PCR positive cases (n=4). Plots A-D represent 4 different patients. Each patient had at least five sequential sera samples collected. Anti-ZIKV IgM was measured by NS1 (Euroimmun; shown as squares) and μ-capture N (Novagnost; shown as circles) ELISAs. Dotted lines represent cut-off values for each assay. Ratios calculated as recommended by the manufacturers. The plots display a unique pattern of ZIKV IgM antibody response over time for each patient.

### Improving the accuracy of the IgG NS1 ELISA

The accuracy of the best performing ZIKV ELISA (i.e. the Euroimmun IgG NS1 ELISA) could be improved by modifying the cut-off used to classify a positive result. The modified cut-off was identified using receiver operator curve (ROC) analyses, defining the point at which the cut-off gave the highest likelihood ratio (33) using all sera from non-ZIKV subjects (n=204) and sera collected ≥7 days among confirmed ZIKV subjects (n=90). Using a cut-off of 1.5, which provided the maximum likelihood ratio (>4.4), the ELISA exhibited an accuracy of 81.0% (previously 77.9%). Sensitivity and specificity were 78.9 and 82.2% respectively (Figure 6; Table S2).

**Figure 6.**
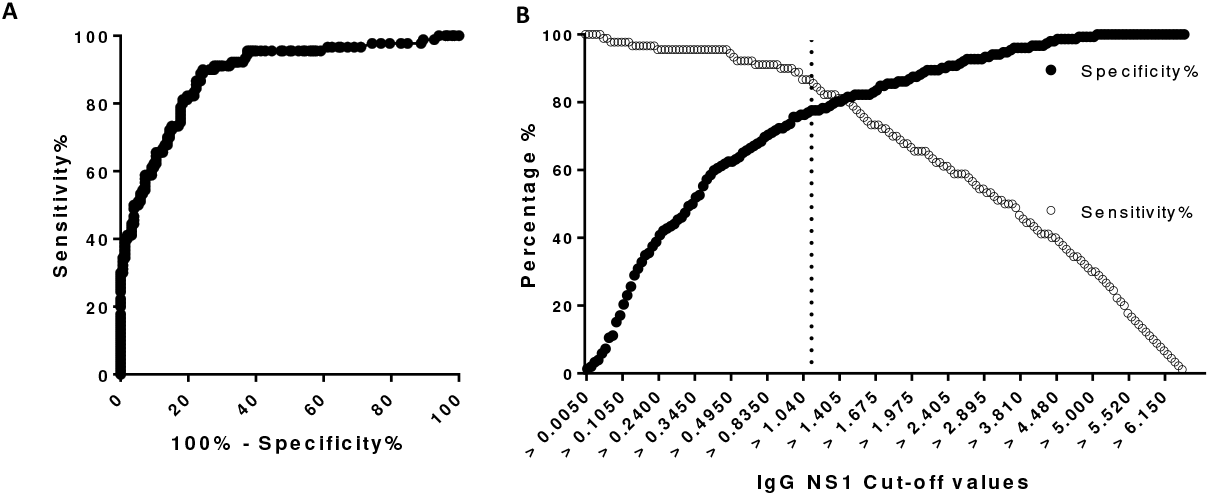
Receiver Operating Characteristic (ROC) curve comparing sensitivity and specificity at different cut-off values for the anti-ZIKV IgG NS1 ELISA (n=294 sera); A) ROC Curve; B) Specificity and Sensitivity at each cut-off. The dotted line in B indicates the cut-off recommended by the manufacturer (ratio of 1.1). The accuracy of IgG NS1 ELISA was 77.9% using the manufacturer’s cut-off. Higher accuracy was observed when the cut-off was increased to 1.5 (where the curves intersect on plot B). Using this cut-off, the ELISA had an accuracy of 81.0%. Sensitivity and specificity were 78.9 and 82.2% respectively.

## Discussion

ZIKV was a viral infection of significant international public health concern that affected over 148 countries during 2015-2019 (5, 34). Pregnant women are still advised not to travel in Brazil and other South American countries. Among people with suspected ZIKV presenting seven or more days from symptom onset serological antibody testing remains the recommended diagnostic approach (32). The majority of ZIKV antibody tests employed in the field have not been validated in their target populations. Despite the ZIKV outbreak in Brazil triggering WHO to declare a public health emergency, there has been no systematic evaluation of the commercial ZIKV antibody assays among Brazilians residents.

The Euroimmun IgG NS1 assay gave the most accurate diagnostic performance among the ELISAs tested. Accuracy could be improved to 81% by modifying the cut-off (from that suggested by the manufacturer). Our results indicate that approximately 1 in 5 subjects are falsely classified by the Euroimmun IgG ELISA when testing a single serum sample. One accepted weakness of employing an IgG based ELISA is that a positive result from a single sample does not discriminate recent from past infection. Akin to other IgG based tests used to diagnose acute infection (35), one option for improving the sensitivity of acute ZIKV diagnosis may be to collect and test serial (paired) samples. During testing of ZIKV PCR positive cases, 95% of paired sera exhibited a rise in antibody levels when measured via the Euroimmun NS1 IgG ELISA (collected seven days apart).

All of the anti-ZIKV IgM ELISAs (Euroimmun NS1, μ-capture and MAC) exhibited lower accuracy (<75%). The ELISAs tended to exhibit low sensitivity for detecting PCR confirmed ZIKV cases, even when serum was collected ≥7 days post symptom onset.

We did not expect the Euroimmun IgG ELISA to give higher sensitivity than the IgM based ELISAs. Over time from symptom onset, the Euroimmun IgG ELISA exhibited more consistent and sustained detection of anti-ZIKV antibody compared to its counter-part anti-ZIKV IgM ELISA. These patterns of IgG and IgM response suggest a secondary immune response to infection (Figures 3 and 7), with an anamnestic boosting of prior immune response most likely due to dengue that cross-reacts in the anti-ZIKV IgG antibody assay (Figure 7). Given our patients presented with their first reported ZIKV infection, their antibody responses suggest they had previously been exposed to a similar virus (i.e. to dengue or another flavivirus). We suggest the prominent IgG response and poor specificity of the anti-Zika IgG assays observed in this study reflect an anamnestic cross-reactive IgG antibody response among local Brazilians who have previously been exposed to other flaviviruses. This warrants further investigation and has implications in the design of both future diagnostic tests and vaccines against ZIKV and DENV in flavivirus exposed populations as it is recognized with dengue.

**Figure 7.**
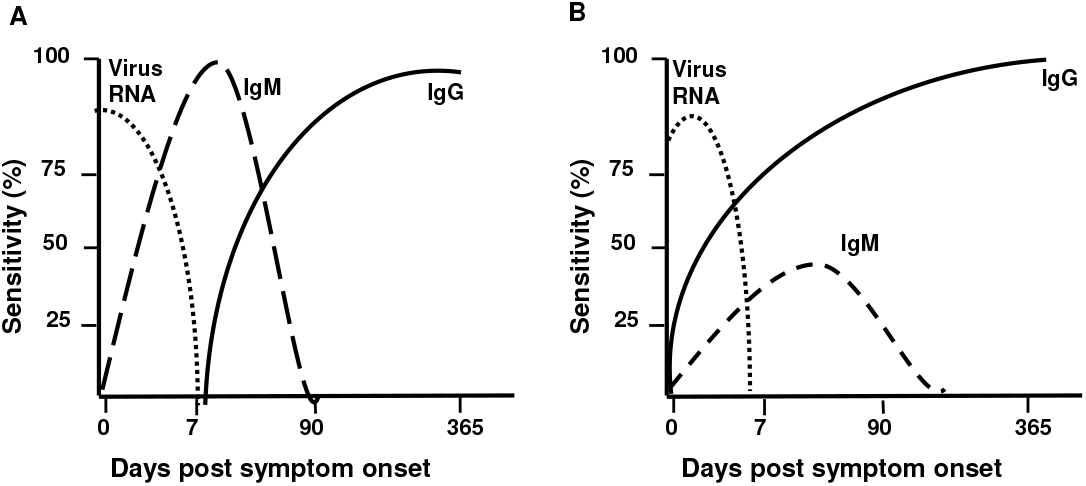
Diagram representing the different patterns of anti-viral IgG and IgM antibody responses and viral RNA detection observed in sera over days from symptom onset among (A) virus naïve and (B) previously exposed individuals. The cartoon exhibits a more prominent IgG response compared to IgM among individuals previously exposed to the virus. In our current study, we observed a more prominent anti-ZIKV IgG compared to IgM response (see Figure 3).

Our findings contrast markedly with published studies conducted using sera from travelers. Such studies largely tested people visiting but not living in ZIKV or other flavivirus exposed countries. These latter studies reported much higher sensitivity (>90%) for the commercial IgM assays (17, 18, 20) and higher specificity for the IgG NS1 assay (>90%) (21). The CDC MAC-ELISA exhibited reduced accuracy and sensitivity when tested among Nicaraguan and Colombian residents compared to “traveler” subjects. Our findings were consistent with this [16,17].

This disparity is likely to reflect the different flavivirus exposure between visitors and local residents. ZIKV positive visitors are likely to experience their first exposure to flavivirus infection. In contrast to residents of Brazil, who are likely to have been exposed to ZIKV and other circulating flaviviruses in the past (e.g. DENV and/or YFV). If past flavivirus exposure triggers an anamnestic antibody response leading to more IgG than IgM production, this could, in part, explain the poor sensitivity of IgM based ZIKV ELISAs seen here, as it has been shown previously for other flaviviruses (36). Further PRNT testing specific for DENV and YFV could be conducted to ascertain prior flavivirus exposure in the specimens evaluated. Our findings indicate that validation of diagnostic assays should be performed in the population it will be used for.

The high rate of DENV IgM assay positivity (43%) among confirmed ZIKV cases was not anticipated (37). The significant correlation between antibody titers for DENV and ZIKV, when tested via IgG or IgM based ELISAs, indicate there is a degree of cross-reaction both ways following DENV and ZIKV infection detected in these ELISAs. These findings highlight the diagnostic challenges ahead as outbreaks of both DENV and ZIKV have been forecast to re-occur in overlapping geographical regions.

Our study findings for the ZIKV diagnostic antibody tests are pertinent to all emerging epidemics, including the current SARS-CoV-2 pandemic. Our results highlight that confirming the accuracy of a diagnostic assay in the target population is imperative to control and manage false positive or negative results across different settings. Such validation should be advocated by governments, national public health agencies and the WHO prior to test deployment. Our results also highlight the need to re-evaluate the accuracy of established tests when a closely-related emergent pathogen is introduced in a region or the population changes. In the case of ZIKV, we recommend the re-evaluation of DENV and YFV assays’ performance in Brazil. However, for newer threats, such as SARS-CoV-2, re-evaluation of Severe Acute Respiratory Syndrome (SARS) coronavirus antibody tests should be considered among Saudi Arabians and other populations with a history of transmission for closely related viruses.

Based on the assays we have assessed, ZIKV PRNTs provide the most accurate assay to diagnose previous exposure to ZIKV among Brazilian residents in samples collected ≥7 days post symptom onset. Performing PRNTs requires specialized training, sophisticated laboratories and the assays are labor intensive; they are therefore unlikely to be widely used for diagnosis outside of reference laboratories (38). As recommended by the WHO, we support their use as a ‘gold standard‘ reference test for flavivirus diagnosis, including ZIKV, if used with an appropriate cutoff to exclude low level cross-reactions (8).

The Euroimmun IgG NS1 ELISA provided the most accurate ELISA test to diagnose previous exposure to ZIKV among Brazilian residents in samples collected ≥7 days post symptom onset. In order to assess whether exposure is acute, we would recommend taking paired samples (7 days apart) and looking for a rise in antibody titers. The best time of collecting for these samples has not been systematically assessed in this study. Based on our observational data, samples collected on days 2 and 9 post symptom onset were associated with the highest fold-changes. However, this was assessed during the first waves of infection with a newly introduced virus and once significant population exposure has occurred interpreting the significance of a positive Zika IgG for acute diagnosis will be even more challenging.

In our study, the ZIKV IgM based ELISAs exhibited poor accuracy (Euroimmun, Novagnost and MAC-ELISA). Similarly, the Panbio DENV ELISAs (particularly IgM) relatively high positivity rate among acute ZIKV cases was a concern. DENV antibody-based serological assays continue to be needed to complement dengue PCR testing. Our findings highlight the need for careful interpretation of existing dengue ELISA results. As more accurate tests are developed, their accuracy in PCR confirmed ZIKV and DENV exposed residents should be comprehensively assessed.

Our study has limitations. Serial samples among ZIKV PCR positive subjects were collected non-systematically as convenience samples. Consequently, we cannot confirm the best time to collect paired samples post symptom onset. Similarly, serial samples were not available for our non-ZIKV subjects, so we were unable to assess the specificity of paired samples testing in the Euroimmun NS1 IgG ELISA. There remains potential for a false positive rise in titers in acutely ill non-ZIKV subjects due to antibody cross-reaction. The ZIKV PCR positive subjects were not tested for DENV by PCR, so we are unable to confirm that the positive DENV IgM results are false. Nevertheless, limited dengue circulation was reported during the study period in the Rio de Janeiro region. Further work is needed.

In conclusion, this is a systematic evaluation of antibody-based ZIKV assays in Brazil. Among ZIKV patients, anti-ZIKV IgG was detected more consistently than IgM, suggesting a secondary antibody response to infection. ZIKV PRNT exhibited the highest accuracy of all assays tested if used with an appropriate cut-off. All ZIKV IgM based ELISAs exhibited low accuracy. The Euroimmun NS1 IgG ELISA exhibited the best ELISA accuracy. Nevertheless, when testing a single serum sample, it misdiagnosed 1 in 5 cases. Testing paired samples via ZIKV IgG based ELISA, may offer a more sensitive method of diagnosing acute ZIKV exposure. Our findings highlight that diagnostic antibody assay use and interpretation needs careful assessment in the target population, particularly when deployed among populations exposed to multiple closely related viruses.

## Acknowledgements

The authors want to thank especially the Flavivirus Laboratory, IOC, Fiocruz including Rita Maria Nogueira, Elaine de Araujo, Simone Sampaio, Carolina Nogueira dos Santos, Celeste, Cintia, Marcus etc. The authors want to thank the Influenza laboratory, IOC, Fiocruz including Marilda Sequeira and the Viral Hepatitis Laboratory including Juliana Gil Melgaço and Paulo Sergio Fonseca de Sousa. RMC thanks the award of the James Porterfield Prize in International Virology.

This work was supported by the United Kingdom Medical Research Council (https://www.mrc.ac.uk/, Grant number MC_PC_15101); the National Institute for Health Research Health Protection Research Unit (NIHR HPRU, http://www.hpruezi.nihr.ac.uk/) in Emerging and Zoonotic Infections at the University of Liverpool, in partnership with Public Health England (PHE) and Liverpool School of Tropical Medicine (LSTM); and the European Union’s Horizon 2020 research and innovation program ZikaPlan under grant agreement No. 734584 (https://ec.europa.eu/programmes/horizon2020/). The views expressed are those of the authors and not necessarily those of the NHS, the NIHR, the Department of Health or Public Health England. The funders had no role in study design, data collection and analysis, decision to publish, or preparation of the manuscript. D B. is partially funded by the European Union’s horizon 2020 research and innovation program (grant agreement no. 734857), ZikAction. P.B. is partially funded by CNPQ 307282/2017-1 and FAPERJ E-26/202.862/2018. LT is supported by the Wellcome Trust (grant number 205228/Z/16/Z),

## References

1. Kindhauser MK, Allen T, Frank V, Santhana RS, Dye C. 2016. Zika: the origin and spread of a mosquito-borne virus. Bull World Health Organ 94:675–686C.

2. Centers for Disease Control and Prevention C. 2018. Zika Travel Information. Traveler’s Health CDC.

3. World Health Organization (WHO). 2019. Zika Epidemiology Update. Zika Virus Disease.

4. (WHO); WHO. 2019. Zika Epidemiology Update. Geneva.

5. Sandra Giron FF, Anne Decoppet, Bernard Cadiou, Thierry Travaglini, Laurence Thirion, Guillaume Duran, Charles Jeannin, Grégory L’Ambert, Gilda Grard, Harold Noël6, Nelly Fournet, Michelle Auzet-Caillaud, Christine Zandotti, Samer Aboukaïs, Pascal Chaud, Saby Guedj, Lakri Hamouda, Xavier Naudot, Anne Ovize, Clément Lazarus, Henriette de Valk, Marie-Claire Paty, Isabelle Leparc-Goffart. 2019. Vector-borne transmission of Zika virus in Europe, southern France, August 2019. Eurosurveillance 24.

6. Aiken ARA, Scott JG, Gomperts R, Trussell J, Worrell M, Aiken CE. 2016. Requests for Abortion in Latin America Related to Concern about Zika Virus Exposure. New England Journal of Medicine 375:396–398.

7. (CDC) CfDCaP. 2017. Guidance for US Laboratories Testing for Zika Virus Infection, July 24, 2017.

8. World Health Organization (WHO). 2016. Laboratory testing for Zika virus infection. Interim guidance World Health Organization. Interim Guidance.

9. Teixeira MCM, C; Barreto, F; Barreto, ML. 2009. Dengue: twenty-five years since reemergence in Brazil. Cad Saude Publica 25

10. Argolo AF, Feres VC, Silveira LA, Oliveira AC, Pereira LA, Junior JB, Braga C, Martelli CM. 2013. Prevalence and incidence of dengue virus and antibody placental transfer during late pregnancy in central Brazil. BMC Infect Dis 13:254.

11. Chiaravalloti-Neto F, da Silva RA, Zini N, da Silva GCD, da Silva NS, Parra MCP, Dibo MR, Estofolete CF, Favaro EA, Dutra KR, Mota MTO, Guimaraes GF, Terzian ACB, Blangiardo M, Nogueira ML. 2019. Seroprevalence for dengue virus in a hyperendemic area and associated socioeconomic and demographic factors using a cross-sectional design and a geostatistical approach, state of Sao Paulo, Brazil. BMC Infect Dis 19:441.

12. Cavalcante KRLJ, & Tauil, Pedro Luiz. 2016. Epidemiological characteristics of yellow fever in Brazil, 2000-2012. Epidemiologia e Serviços de Saúde 25:11–20.

13. Organization. WH. 2018. Yellow fever—Brazil. Disease outbreak news. Emergency preparecness response Report.. Geneva, Switzerland: World Health Organization;.

14. Charrel RL-G, I; Pas, S; Lamballerie, X; Koopmans, M. & Reusken, C.. 2016. Background Review for diagnostic test development for Zika virus infection. Bulletin of the World Health Organization 94:574.

15. Murtagh MM. 2017. Zika Virus Infection Diagnostics Landscape. The International Diagnostics Centre

16. Michelson Y, Lustig Y, Avivi S, Schwartz E, Danielli A. 2019. Highly Sensitive and Specific Zika Virus Serological Assays Using a Magnetic Modulation Biosensing System. J Infect Dis 219:1035–1043.

17. Huzly D, Hanselmann I, Schmidt-Chanasit J, Panning M. 2016. High specificity of a novel Zika virus ELISA in European patients after exposure to different flaviviruses. Euro Surveill 21.

18. Steinhagen K, Probst C, Radzimski C, Schmidt-Chanasit J, Emmerich P, van Esbroeck M, Schinkel J, Grobusch MP, Goorhuis A, Warnecke JM, Lattwein E, Komorowski L, Deerberg A, Saschenbrecker S, Stocker W, Schlumberger W. 2016. Serodiagnosis of Zika virus (ZIKV) infections by a novel NS1-based ELISA devoid of cross-reactivity with dengue virus antibodies: a multicohort study of assay performance, 2015 to 2016. Euro Surveill 21.

19. Van Esbroeck M, Meersman K, Michiels J, Arien KK, Van den Bossche D. 2016. Letter to the editor: Specificity of Zika virus ELISA: interference with malaria. Euro Surveill 21.

20. L’Huillier AG, Hamid-Allie A, Kristjanson E, Papageorgiou L, Hung S, Wong CF, Stein DR, Olsha R, Goneau LW, Dimitrova K, Drebot M, Safronetz D, Gubbay JB. 2017. Evaluation of Euroimmun Anti-Zika Virus IgM and IgG Enzyme-Linked Immunosorbent Assays for Zika Virus Serologic Testing. J Clin Microbiol 55:2462–2471.

21. Safronetz D, Sloan A, Stein DR, Mendoza E, Barairo N, Ranadheera C, Scharikow L, Holloway K, Robinson A, Traykova-Andonova M, Makowski K, Dimitrova K, Giles E, Hiebert J, Mogk R, Beddome S, Drebot M. 2017. Evaluation of 5 Commercially Available Zika Virus Immunoassays. Emerg Infect Dis 23:1577–1580.

22. Balmaseda A, Zambrana JV, Collado D, Garcia N, Saborio S, Elizondo D, Mercado JC, Gonzalez K, Cerpas C, Nunez A, Corti D, Waggoner JJ, Kuan G, Burger-Calderon R, Harris E. 2018. Comparison of Four Serological Methods and Two Reverse Transcription-PCR Assays for Diagnosis and Surveillance of Zika Virus Infection. J Clin Microbiol 56.

23. Granger D, Hilgart H, Misner L, Christensen J, Bistodeau S, Palm J, Strain AK, Konstantinovski M, Liu D, Tran A, Theel ES. 2017. Serologic Testing for Zika Virus: Comparison of Three Zika Virus IgM-Screening Enzyme-Linked Immunosorbent Assays and Initial Laboratory Experiences. J Clin Microbiol 55:2127–2136.

24. Brasil P, Calvet GA, Siqueira AM, Wakimoto M, de Sequeira PC, Nobre A, Quintana Mde S, Mendonca MC, Lupi O, de Souza RV, Romero C, Zogbi H, Bressan Cda S, Alves SS, Lourenco-de-Oliveira R, Nogueira RM, Carvalho MS, de Filippis AM, Jaenisch T. 2016. Zika Virus Outbreak in Rio de Janeiro, Brazil: Clinical Characterization, Epidemiological and Virological Aspects. PLoS Negl Trop Dis 10:e0004636.

25. Centers for Disease Control and Prevention C. 2016. Zika MAC-ELISA: For Use Under an Emergency Use Authorization Only - Instructions for Use.

26. Russell PK, Nisalak A, Sukhavachana P, Vivona S. 1967. A plaque reduction test for dengue virus neutralizing antibodies. J Immunol 99:285–90.

27. Castanha PM, Cordeiro MT, Martelli CM, Souza WV, Marques ET, Jr., Braga C. 2013. Force of infection of dengue serotypes in a population-based study in the northeast of Brazil. Epidemiol Infect 141:1080–8.

28. Castanha PMS, Souza WV, Braga C, Araujo TVB, Ximenes RAA, Albuquerque M, Montarroyos UR, Miranda-Filho DB, Cordeiro MT, Dhalia R, Marques ETA, Jr., Rodrigues LC, Martelli CMT, Microcephaly Epidemic Research G. 2019. Perinatal analyses of Zika- and dengue virus-specific neutralizing antibodies: A microcephaly case-control study in an area of high dengue endemicity in Brazil. PLoS Negl Trop Dis 13:e0007246.

29. Lanciotti RS, Kosoy OL, Laven JJ, Velez JO, Lambert AJ, Johnson AJ, Stanfield SM, Duffy MR. 2008. Genetic and serologic properties of Zika virus associated with an epidemic, Yap State, Micronesia, 2007. Emerg Infect Dis 14:1232–9.

30. Santiago GA, Vergne E, Quiles Y, Cosme J, Vazquez J, Medina JF, Medina F, Colon C, Margolis H, Munoz-Jordan JL. 2013. Analytical and clinical performance of the CDC real time RT-PCR assay for detection and typing of dengue virus. PLoS Negl Trop Dis 7:e2311.

31. Bossuyt PM, Reitsma JB, Bruns DE, Gatsonis CA, Glasziou PP, Irwig L, Lijmer JG, Moher D, Rennie D, de Vet HC, Kressel HY, Rifai N, Golub RM, Altman DG, Hooft L, Korevaar DA, Cohen JF, Group S. 2015. STARD 2015: an updated list of essential items for reporting diagnostic accuracy studies. BMJ 351:h5527.

32. World Health Organization. 2016. Laboratory testing for Zika virus infection. Interim guidance 23 March 2016. http://apps.who.int/iris/bitstream/10665/204671/1/WHO_ZIKV_LAB_16.1_eng.pdf?ua=1. Accessed 08/09/2016.

33. Hajian-Tilaki K. 2013. Receiver Operating Characteristic (ROC) Curve Analysis for Medical Diagnostic Test Evaluation. Caspian J Intern Med 4:627–35.

34. Organization WH. 2018. Zika virus (ZIKV) classification table data as of 15 February 2018 [Internet]. Geneva; World Health Organization.

35. Hang VT, Nguyet NM, Trung DT, Tricou V, Yoksan S, Dung NM, Van Ngoc T, Hien TT, Farrar J, Wills B, Simmons CP. 2009. Diagnostic accuracy of NS1 ELISA and lateral flow rapid tests for dengue sensitivity, specificity and relationship to viraemia and antibody responses. PLoS Negl Trop Dis 3:e360.

36. Solomon T, Thao LT, Dung NM, Kneen R, Hung NT, Nisalak A, Vaughn DW, Farrar J, Hien TT, White NJ, Cardosa MJ. 1998. Rapid diagnosis of Japanese encephalitis by using an immunoglobulin M dot enzyme immunoassay. J Clin Microbiol 36:2030–4.

37. Honorio NA, Nogueira RM, Codeco CT, Carvalho MS, Cruz OG, Magalhaes Mde A, de Araujo JM, de Araujo ES, Gomes MQ, Pinheiro LS, da Silva Pinel C, Lourenco-de-Oliveira R. 2009. Spatial evaluation and modeling of Dengue seroprevalence and vector density in Rio de Janeiro, Brazil. PLoS Negl Trop Dis 3:e545.

38. Thomas SJ, Nisalak A, Anderson KB, Libraty DH, Kalayanarooj S, Vaughn DW, Putnak R, Gibbons RV, Jarman R, Endy TP. 2009. Dengue plaque reduction neutralization test (PRNT) in primary and secondary dengue virus infections: How alterations in assay conditions impact performance. Am J Trop Med Hyg 81:825–33.

